# Myosin 7b is a regulatory long noncoding RNA (lncMYH7b) in the human heart

**DOI:** 10.1101/2020.09.06.285189

**Authors:** Lindsey J. Broadwell, Michael J. Smallegan, Kevin M. Rigby, Jose S. Navarro-Arriola, Rusty L. Montgomery, John L. Rinn, Leslie A. Leinwand

**Affiliations:** Dept of Biochemistry, CU Boulder; Biofrontiers Institute, CU Boulder; Dept of Molecular, Cellular, and Developmental Biology, CU Boulder; miRagen, Boulder, CO

**Keywords:** Heart failure, myosin heavy chain (MyHC), myosin heavy chain 7b (MYH7b), alternative, splicing, long noncoding RNA (lncRNA), gene regulation

## Abstract

Myosin heavy chain 7b (MYH7b) is an ancient member of the myosin heavy chain motor protein family that is expressed in striated muscles. In mammalian cardiac muscle, MYH7b RNA is expressed along with two other myosin heavy chains, β-myosin heavy chain (β-MyHC) and α-myosin heavy chain (a-MyHC). However, unlike β-MyHC and α-MyHC which are maintained in a careful balance at the protein level, the MYH7b locus does not produce protein in the heart due to a post-transcriptional exon-skipping mechanism that occurs in a tissue specific manner. Whether this locus has a role in the heart beyond producing its intronic microRNA, miR-499, was unclear. Using cardiomyocytes derived from human induced pluripotent stem cells as a model system, we have found that the non-coding exon skipped RNA (lncMYH7b) affects the transcriptional landscape of the human heart, independent of miR-499. Specifically, lncMYH7b regulates the ratio of β-MyHC to α-MyHC, which is crucial for heart function. This regulation is likely achieved through control of members of the TEA domain transcription factor family (TEAD1 and TEAD3). Therefore, we conclude that this ancient gene has been repurposed by alternative splicing to produce a regulatory long-noncoding RNA in the human heart that affects the myosin composition of the heart.

## INTRODUCTION

The myosin family of motor proteins that drives striated muscle contraction consists of 10 genes with distinct functions(1). Three of these genes are expressed in mammalian hearts (*MYH6, MYH7*, and *MYH7b*). *MYH6* (α-MyHC) and *MYH7* (β-MyHC) are the major sarcomeric myosin proteins expressed in mammalian hearts. In humans, >90% of the heart’s myosin protein composition is comprised of β-MyHC with the remaining <10% being α-MyHC and the two are antithetically regulated (1–5). However, various conditions can shift their relative proportions. The finely-tuned balance of these two motors is critical for proper cardiac function since they have very different enzymatic properties which determine the contractile velocity of the muscle (6). It has been well-established that disturbing the β-/α-MyHC ratio leads to compromised contractility in cardiomyocytes; in human heart failure, there is a shift to ~100% β-MyHC and α-MyHC becomes undetectable (2–4, 7). As β-MyHC is the slower, more efficient motor, this shift in expression is thought to be an initial compensatory mechanism to preserve energy in the failing heart. Heart failure patients that show functional improvement upon treatment with β-adrenergic receptor blockers have an increase in α-MyHC expression (4). This suggests that maintenance of the β-MyHC/α-MyHC ratio is fundamental to proper cardiac function.

While *MYH6* and *MYH7* have been studied for decades, *MYH7b* was not identified until the sequencing of the human genome and was initially annotated as a sarcomeric myosin based on sequence alignments with other known family members (8). Based on phylogenetic analysis, *MYH7b* was identified as an ancient myosin, indicating that *MYH7b* was present before the gene duplication events that led to the other sarcomeric myosins (8). The human *MYH7b* gene is located on chromosome 20, separate from the two canonical sarcomeric myosin clusters on chromosomes 14 (cardiac myosins) and 17 (skeletal muscle myosins), supporting the idea that *MYH7b* may have a specialized role in muscle biology (1, 8). In certain species, including snakes and birds, *MYH7b* is expressed as a typical sarcomeric myosin protein in heart and skeletal muscle (9). However, *MYH7b* has a unique expression pattern in mammals: the encoded protein is restricted to specialized muscles and non-muscle tissues (10–12). Furthermore, while *MYH7b* RNA is expressed in mammalian cardiac and skeletal muscle, an alternative splicing event skips an exon, introducing a premature termination codon (PTC) that prevents full-length protein expression (Fig 1A) (10). This raises the question of why the MYH7b RNA expression has been conserved in mammals. One obvious hypothesis is that *MYH7b* transcription is preserved in mammalian heart and skeletal muscle in order to maintain the expression of an *MYH7b* intronic microRNA, miR-499, in those tissues (10). However, miR-499 knockout mice have no discernible cardiac or skeletal muscle phenotype (13). It’s possible that the species-specific regulation of myosin isoforms complicates the interpretation of this mouse model and that miR-499 could play an important role in human muscle (6, 14). Another hypothesis is that a heretofore unknown functional molecule is expressed from the *MYH7b* gene, causing its conserved expression in sarcomeric muscle tissue. In this report, we performed a thorough dissection of this locus in order to discern the role of *MYH7b* gene expression in human cardiomyocytes and found a novel long noncoding RNA (lncMYH7b) with roles in cardiac gene expression.

**Figure 1.**
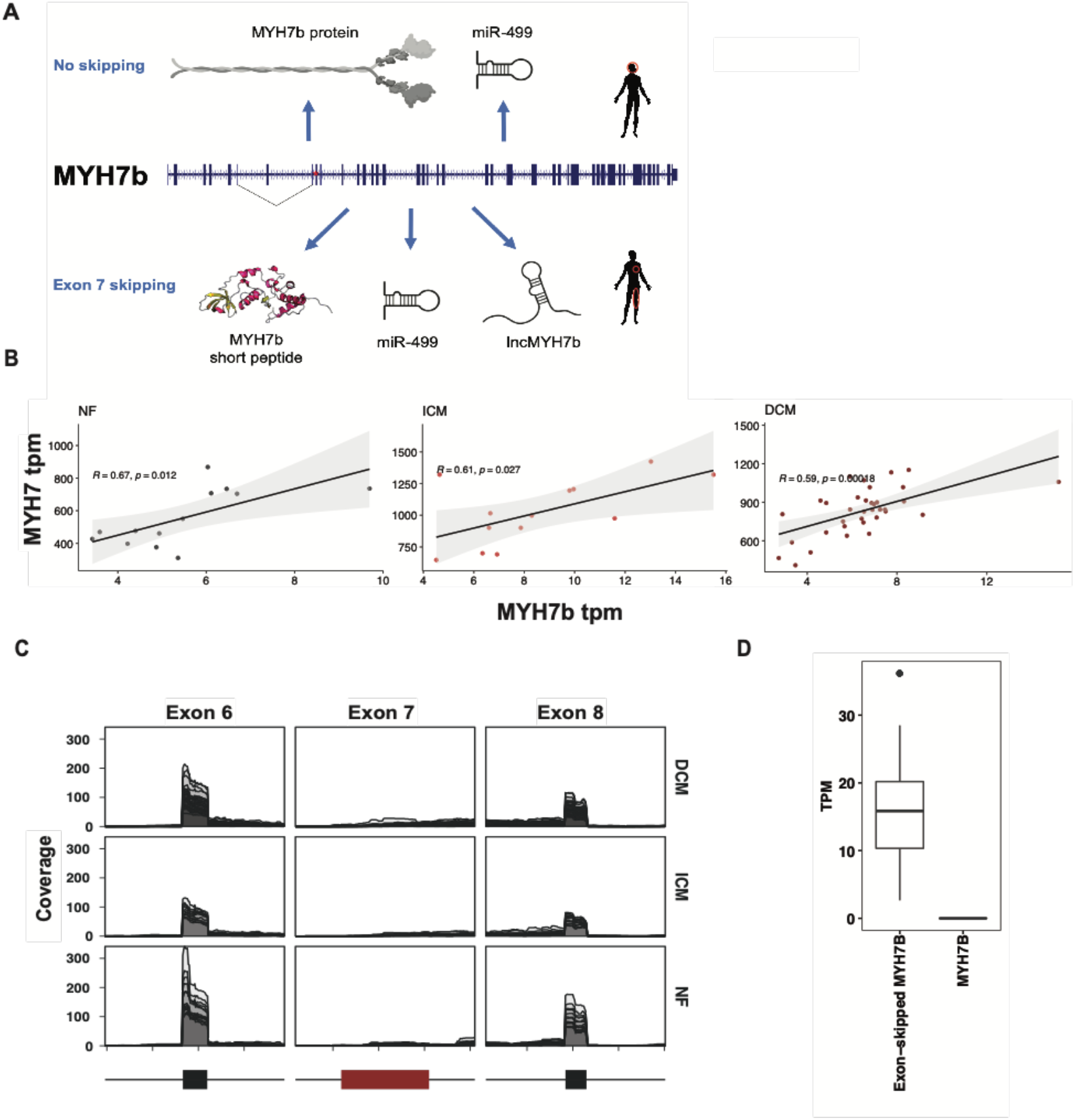
*MYH7b* is alternatively spliced in the heart and correlates with β-MyHC in heart disease. **A)** *MYH7b* is expressed as a full-length sarcomeric myosin protein in the human brain and inner ear, but is alternatively spliced in the heart and skeletal muscle to become non-coding. A putative short peptide (MYH7b_sp) could be produced from the exon-skipped transcript, and a microRNA (miR-499) is encoded regardless of splicing. **B)** The expression of exon 7-skipped MYH7b correlates with that of β-MyHC in non-failing (NF), dilated cardiomyopathy (DCM), and ischemic cardiomyopathy (ICM). Data pulled from GSE116250. **C)** Splicing analysis of MYH7b in 64 human hearts shows that exon 7 is almost nonexistent in the human heart. **D)** Quantification of exon-skipping in human hearts.

## MATERIALS AND METHODS

### Cell Culture

The WTC-11 iPSC line was used for all RNA-seq and FISH experiments. A small molecule differentiation protocol was followed (15). Wnt signaling was activated for 48 hrs using Chiron 99021 (GSK3 inhibitor) at 5 μM in RPMI base media with BSA and L-ascorbic acid. Wnt was then inhibited for 48 hrs using Wnt-C59 at 2 μM in the same base media. The media was then switched to a B27 supplement with insulin in RPMI for the remainder of the culture time. All cells were matured for 35 days before treatment with ASO or virus.

iCell Cardiomyocytes were purchased from Cellular Dynamics International (CMC-100-010-001), cultured to manufacturer’s protocols, and used for all qPCR experiments. qPCR probers were purchased from Applied Biosystems and performed according to manufacturer’s protocols.

### ASO Treatment

Matured cardiomyocytes were passively treated with either a scrambled (control) ASO or one targeting MYH7b at 2 μM for 96 hrs. ASO was refreshed with media changes, every 48 hrs. ASO was manufactured and verified by miRagen (sequence and chemistry proprietary). For the kinetic study, iCell Cardiomyocytes were treated with 10 uM ASO targeting MYH7b for indicated timepoints. ASO was refreshed with media changes every 24 hours and RNA was collected at indicated timepoints.

### FISH Probe Design

Probes were designed against the human MYH7b using the LGC Biosearch Technologies’ Stellaris probe designer (Stellaris Probe Designer v. 4.2). As there was a concern for cross-reaction with both β-MyHC and α-MyHC due to high sequence conservation in the myosin II family, these probes were ranked according to how many mismatches they had with the other two transcripts. The top 29 were ordered from Integrated DNA Technologies and labeled with ATTO-550 dye using terminal deoxynucleotidyl transferase (TdT) as described in Gaspar et al 2017. Briefly, TdT was incubated with 5-Propargylamino-ddUTP-ATTO-550 and an equimolar mix of the 29 probes at 37°C for 16 hrs, refreshing the TdT at 8 hrs. The labeled probes were then purified through ethanol precipitation.

Matured cardiomyocytes were replated on plasma cleaned coverslips coated in a GelTrex cushion. 48 hrs after, the cardiomyocytes were washed three times with PBS, then fixed with 4% PFA for 10 min at room temperature. After washing again, coverslips were transferred to 70% ethanol and kept at 4°C for at least 1 hr. The 70% ethanol was then removed and 1 mL Stellaris RNA FISH Wash Buffer A (Biosearch Technologies Cat# SMF-WA1-60) with 10% deionized formamide for 5 min at room temperature. A humidifying chamber was assembled in a 15 cm tissue culture plate using a wet paper towel covered in Parafilm. For each coverslip, 2 μL of the TdT-labeled probes was added to 98 μL of Stellaris RNA FISH Hybridization Buffer (Biosearch Technologies Cat# SMF-HB1-10); 100 μL of this mixture was dotted onto the Parafilm in the humidifying chamber. The chamber was sealed with Parafilm then left in the dark at 37 °C overnight. Coverslips were then transferred back into a 12-well culture plate and washed twice with 1 mL Buffer A for 30 min at 37°C, then in 1 mL Stellaris RNA FISH Wash Buffer B (Biosearch Technologies Cat# SMF-WB1-20) with Hoescht stain at 1:10,000 dilution for 5 min at RT. Coverslips were then mounted using Vectashield anti-fade mounting medium and sealed with clear nail polish. Cells were imaged with a DeltaVision confocal microscope and images were deconvolved using the Softworx software. Localization was determined using Imaris to detect all dots, then only dots within nuclei and taking the ratio of those two measurements; 100 nuclei were analyzed across 8 field of views in two independent experiments.

### Adenovirus production

Adenovirus for recombinant expression of MYH7b_sp was produced as has been described before using the pAdEasy system (16).

### Whole cell lysates, myosin separating gels and western blotting

All tissue samples were powdered with a BioPulverizer then nutated in RIPA buffer (140 mM NaCl, 50 mM Tris, 0.5 mM EGTA, 1 mM EDTA, 1% Triton X-100, 0.1% SDS, 0.5% deoxycholate) for 45 min at 4°C then spun at 15,000 rpm for 15 min to pellet the insoluble fraction. Cell lysates were prepared in the same manner. Lysates were quantified using a BioRad DC Assay. To determine the myosin isoform % in control vs MYH7b KD samples, we followed the protocol in Warren and Greaser[4]. Briefly, 4 μg of each sample was run on a 6% polyacrylamide-N,N0-diallyltartardiamide (DATD) gel and stained with Sypro. Relative percentages of α-MyHC and β-MyHC were determined for each lane using ImageQuant. For western blotting, 40 μg of protein was run on a 12% SDS-PAGE gel and transferred to a PVDF membrane. An anti-MYH7b antibody was used at 1:100 (11, 17).

### RNA isolation

Total RNA was isolated using Tri reagent. Samples were suspended in Tri reagent, then total RNA was organically extracted using chloroform. RNA was then alcohol precipitated with isopropanol and resuspended in sterile water.

### RNA-sequencing and analysis

RNA samples were submitted to Novogene for library preparation, by PolyA selection, and sequencing. All samples had a sequencing depth of at least 20 million 150 bp paired end reads. The reads were mapped and extensive quality control was performed using the nf-core/rnaseq v1.4.2 pipeline. Reads were mapped to hg38 and gene quantification used Gencode v34. Differential expression analysis was performed using DESeq2 in R v3.5.1. Functional enrichment was performed with ClusterProfiler in R. All scripts are available on GitHub at: https://github.com/libr8211/lncMYH7b

RNA-seq reads for the human heart data were retrieved from GEO (accession number GSE116250). Since the exon-skipped transcript is not present in the Gencode v34 annotation, we appended the specific exon-skipped transcript entry for MYH7b to the Gencode human annotation v34 and ran the same bioinformatic analysis as above to quantify the amount of exon skipped vs full length transcript. Details are available in the same Github repository as above.

## RESULTS

### MYH7b and β-MyHC expression are positively coregulated

The human MYH7b locus is complex. In certain tissues, it encodes a typical striated muscle myosin mRNA and full-length protein. In tissues where exon 7 is skipped, the locus could theoretically produce both a peptide originating from the short open reading frame preceding the PTC in exon 9 (MYH7b_sp) and the exon 7-skipped long-noncoding RNA itself (lncMYH7b, Fig. 1A). In both contexts, the microRNA miR-499 is encoded in intron 19. We analyzed RNA-sequencing (RNA-seq) data from 64 healthy and diseased human heart samples and found that exon 7 (the skipped exon) of the *MYH7b* RNA is essentially undetectable regardless of disease state, which is consistent with our previous observation that *MYH7b* has a tissue-specific alternative splicing pattern (Fig 1B-C) (10). This transcriptomic analysis suggests that the full-length MYH7b protein is not present in the human heart. This observation is also consistent with our previous results showing that forced expression of full-length MYH7b protein in the mouse heart results in cardiac dilation and dysfunction, despite roles the full-length protein is known to have in other tissues (12, 17, 18).

β-MyHC induction, which shifts the β-MyHC/α-MyHC ratio, is a hallmark of chronic heart disease (2–4). Intriguingly, we observed that *MYH7b* RNA levels correlate with β-MyHC expression, including the known increase in β-MyHC expression in diseased human hearts (Fig 1D, S1). This raised the possibility that there is a regulatory relationship between the *MYH7b* locus and β-MyHC expression. Although full-length MYH7b protein and mRNA are essentially absent in the human heart, there are several other molecules originating from the *MYH7b* gene locus, as mentioned above. The presence of these different molecules prompted further investigation into the locus as a whole.

### miR-499 does not regulate β-MyHC expression

One hypothesis is that the MYH7b locus regulates β-MyHC through the miR-499 encoded in intron 19 of the *MYH7b* pre-mRNA. There is no evidence to suggest that miR-499 regulates the expression of either α- or β-MyHC, and the miR-499 knockout mouse still induces β-MyHC in the heart under conditions of hypothyroidism (10, 13, 19). Other microRNAs, such as miR-208a and miR-208b encoded by the α-MyHC and β-MyHC genes, respectively, have been shown to have effects on the β-MyHC/α-MyHC ratio in mouse models (13). However, the adult mouse cardiac β-MyHC/α-MyHC ratio is opposite to the human one (5%:95% compared to 9O%:IO%), demonstrating species-specific regulation (6, 14). Therefore, it was important to investigate the β-MyHC response to changes in miR-499 activity in a human context. We employed a cellular model of human induced pluripotent stem cells differentiated to cardiomyocytes (hiPS-CMs), which we treated with an anti-miR that targets miR-499 specifically. We observed no changes in *MYH7b* or β-MyHC levels (Fig. 2). Therefore, we conclude that miR-499 does not regulate β-MyHC expression in human cardiomyocytes.

**Figure 2.**
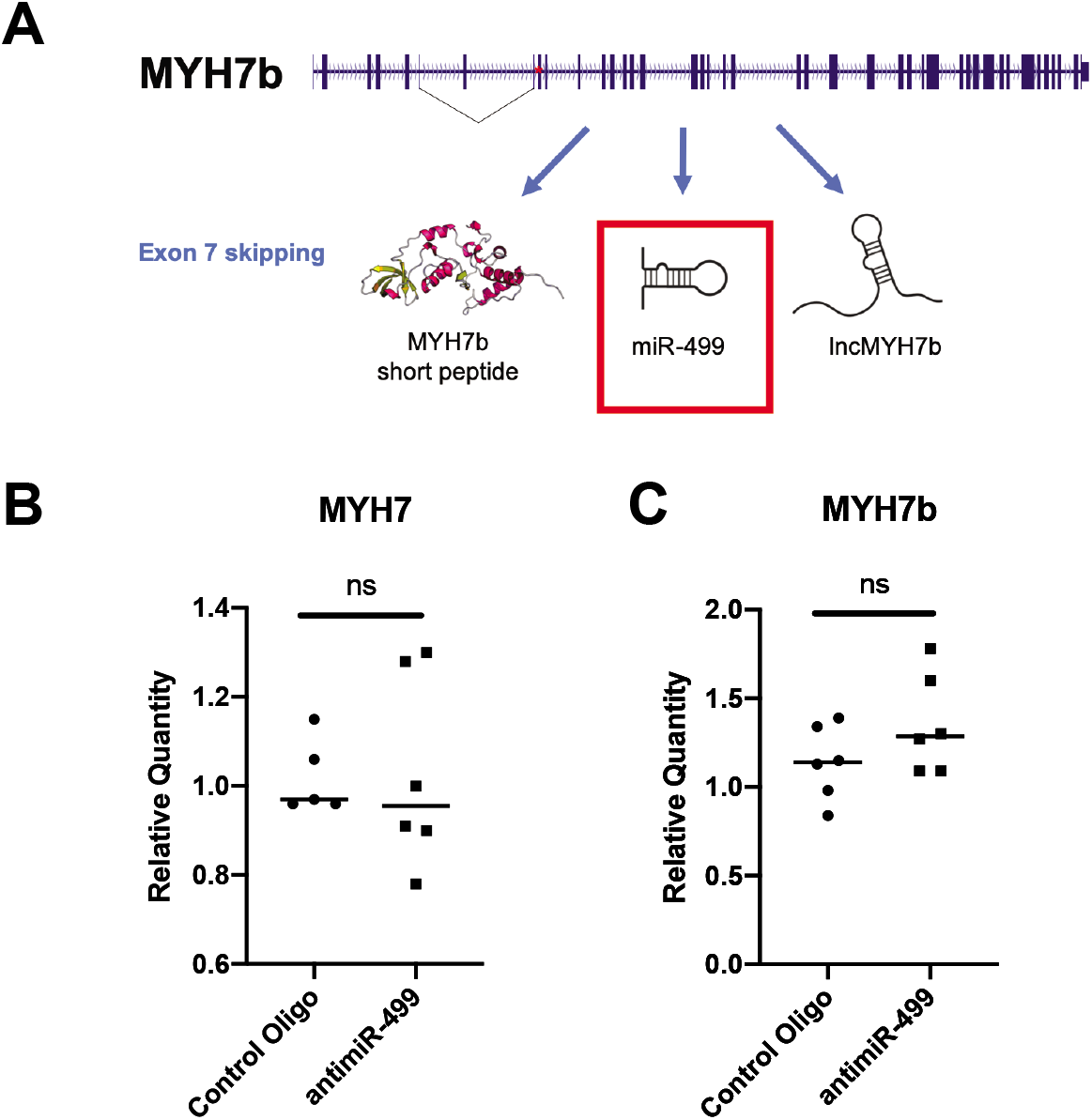
miR-499 does not affect the expression of either MYH7b or β-MyHC. **A)** miR-499 is still produced upon skipping of exon 7 and could have a role in β-MyHC regulation. **B)** An anti-miR targeted to miR-499 does not change the expression levels of MYH7b or β-MyHC.

### MYH7b short peptide is not expressed in human hearts

The elimination of miR-499 as a regulator of the β-MyHC/α-MyHC ratio left both a putative short peptide and the lncMYH7b itself to investigate (Fig. 3A). The *MYH7b* locus produces fUll-length protein in specialized muscles, brain, and inner ear tissues and therefore encodes all of the essential signals for translation (9). We previously reported increased exon 7-skipped *MYH7b* RNA levels in C2C12 mouse myotubes after blocking translation with cycloheximide, indicating that the exon-skipped RNA is either translated or shunted to the nonsense-mediated decay (NMD) pathway (10). Using fluorescent *in situ* hybridization (FISH), we determined that MYH7b RNA is localized to the cytoplasm in hiPS-CMs, consistent with translation (Fig 3B-C). Furthermore, we saw very few transcripts along the nuclear envelope, which is where mRNAs undergoing NMD typically localize (Fig 3B) (20). This suggests that not all exon 7-skipped *MYH7b* RNA is degraded by NMD. There is also precedent for small peptides being produced from “non-coding” RNAs which affect muscle contraction (21, 22). Based on this, we next investigated whether the short open reading frame prior to the PTC in exon 9 is translated and, if so, whether the short peptide (MYH7b_sp) could regulate the β-MyHC/α-MyHC ratio in human cardiomyocytes. The putative MYH7b_sp is predicted to be 206 amino acids in length and approximately 27 kDa. It is possible that MYH7b_sp has escaped detection in previous studies simply because full-length MyHCs are approximately 250 kDa, and a 27 kDa MyHC has not been described. Therefore, we used our anti-MYH7b antibody, which is directed against an epitope that should present in both the full-length MYH7b (as shown in Fig 3D, mouse cerebellum) and the putative MYH7b_sp to probe for MYH7b_sp expression (11, 17). As a positive control, we infected C2C12 cells with an adenovirus containing only the MYH7b_sp open reading frame and detected a protein of the right size and immunoreactivity (Fig 3D). We probed human heart tissue from dilated cardiomyopathy (DCM) patients, where MYH7b transcripts are easily detectable (Fig 1B), but did not detect it via western blotting (Fig 3D). In case MYH7b_sp exists at levels below the detection limit of a western blot but does have a regulatory role, we forced expression of the MYH7b_sp using adenovirus infection of hiPS-CMs and performed RNA-seq (see Fig. 5B). We found no difference in the β-MyHC/α-MyHC ratio at the RNA level, and many differentially expressed genes were chaperones, which is both consistent with an adenoviral infection and production of a foreign peptide (Fig S2) (23). Thus, we conclude that MYH7b_sp is not expressed at detectable levels in human cardiomyocytes and has no effect on the expression of α- or β-MyHC, even after forced expression.

**Figure 3.**
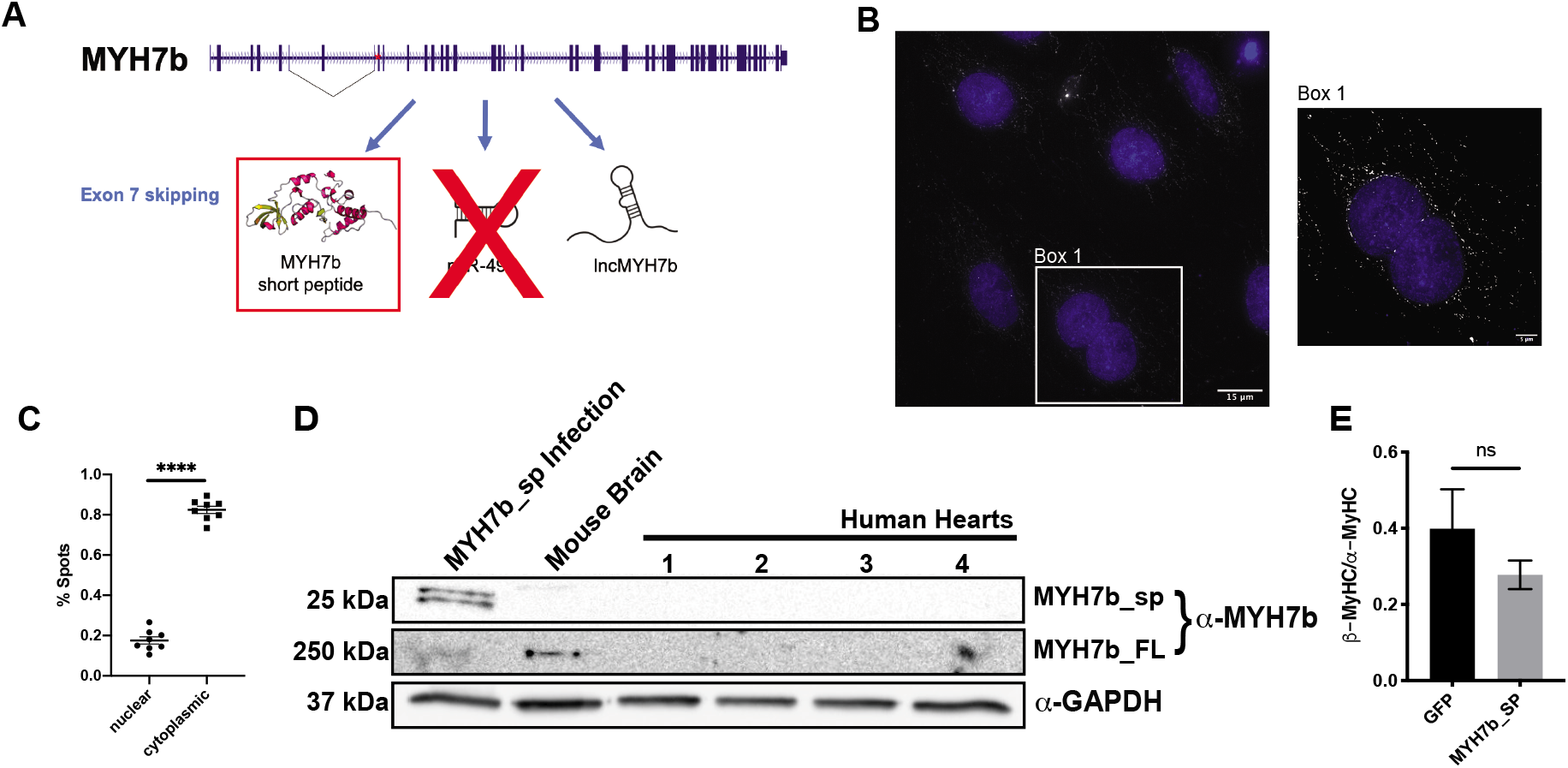
MYH7b_sp does not affect the β-/α-MyHC ratio and is undetectable in human hearts. **A)** miR-499 has been eliminated as the active molecule involved in MyHC regulation in the heart, however MYH7b_sp could have a regulatory role. **B)** Single molecule FISH shows that MYH7b RNA is largely is localized in the cytoplasm, but not along the nuclear envelope where NMD largely occurs. **C)** Quantification of MYH7b RNA localization using FISH. **D)** Western blotting of failing human hearts cannot detect MYH7b_sp using a custom MYH7b antibody that does detect full-length MYH7b in mouse cerebellum. **E)** RNA-seq of MYH7b_sp overexpression shows no significant change in the β-MyHC/α-MyHC ratio.

### lncMYH7b has a role in β-MyHC regulation in human cardiomyocytes

Having ruled out the possible regulatory roles of miR-499 and MYH7b_sp in β-MyHC expression, we hypothesized that the lncMYH7b transcript itself regulates β-MyHC. To test this, we knocked down MYH7b RNA with locked nucleic acid antisense oligonucleotides (ASOs) and measured the kinetics of lncMYH7b and β-MyHC expression in iPS-CMs. .Upon treatment with 10μM ASO, MYH7b RNA was knocked down ~12-fold by 48 hr, which was maintained through 7 days (Fig 4A). Importantly, we observed decreased β-MyHC expression only after loss of MYH7b RNA (Fig 4A), supporting an upstream role for the exon-skipped MYH7b RNA in regulating β-MyHC expression. To further support the idea that lncMYH7b is regulating β-MyHC/α-MyHC ratios independently of miR-499, we measured the change in myomiR levels after lncMYH7b knockdown (KD). We observed no change in miR-499 levels, nor in the other two myomiRs encoded by α-MyHC and β-MyHC (miR-208a and miR-208b, respectively) upon treatment with the MYH7b ASO (Fig 4B).

**Figure 4.**
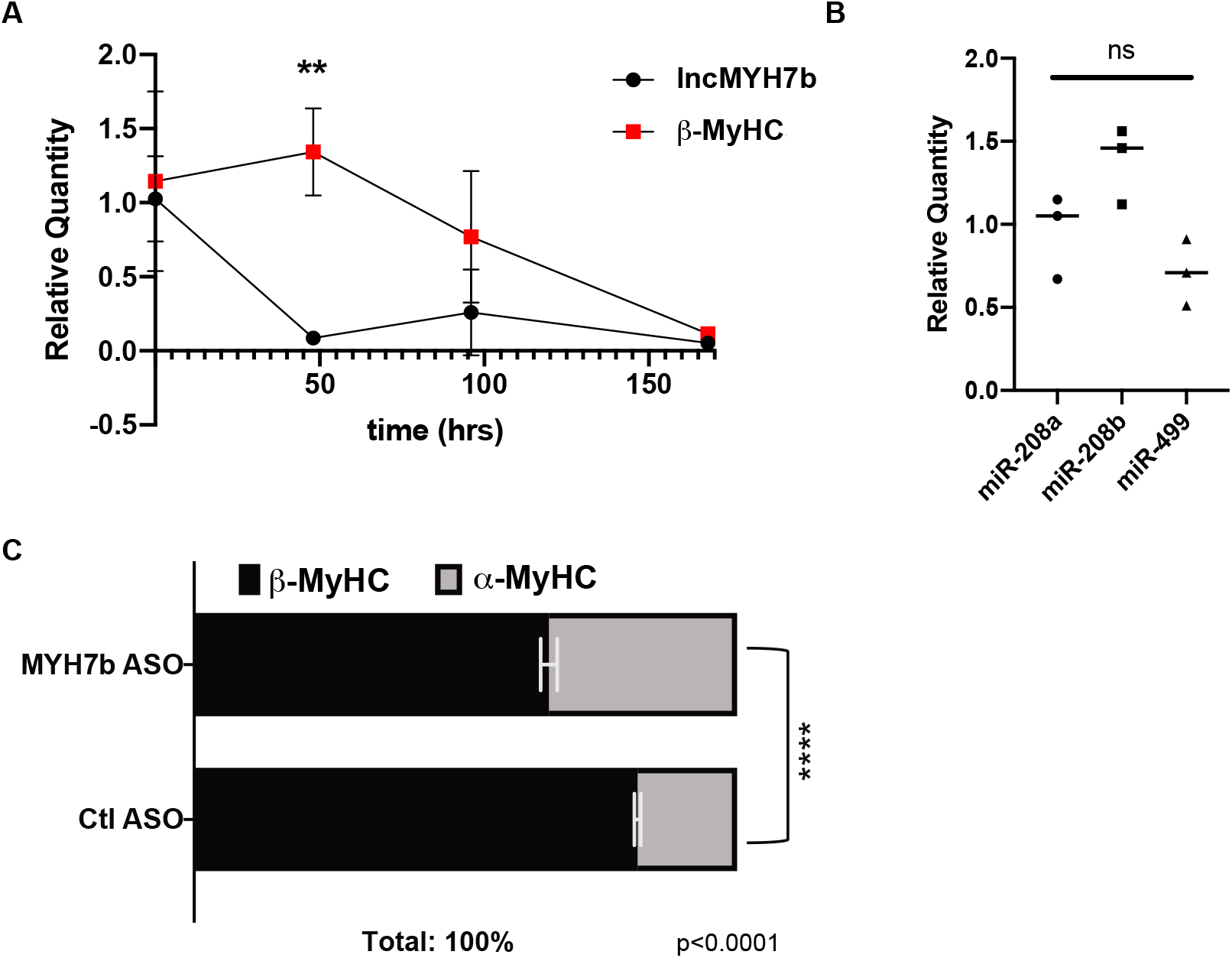
lncMYH7b controls β-MyHC expression at both the RNA and protein level. **A)** MYH7b RNA decreases at least 48 hrs before there is a change in β-MyHC RNA levels are reduced. **B)** ASOs targeted against MYH7b RNA do not affect myomiR levels. **C)** A myosin separating gel shows a reduction in the β-/α-MyHC ratio at the protein level upon lncMYH7b KD in hiPS-CMs.

Next, we needed to confirm that the change observed in β-MyHC RNA upon MYH7b KD was recapitulated at the protein level. As independent validation, we differentiated hiPS-CMs from a different iPSC line and used a different ASO sequence. A myosin separating gel showed a significant decrease of the β-MyHC/α-MyHC ratio upon KD of MYH7b RNA (Fig 4C). Together, these data confirm the role of lncMYH7b in regulating the β-MyHC/α-MyHC ratio in human cardiomyocytes.

In order to elucidate the potential regulatory impacts of lncMYH7b in a more global and unbiased manner, we performed RNA-seq on MYH7b ASO-treated hiPS-CMs. We used the publicly available NF-core/RNAseq pipeline to analyze differential gene expression (24). Differential expression analysis revealed a large bias towards downregulation (535 downregulated vs 224 upregulated genes) with very little overlap with our MYH7b_sp dataset (Fig. 5B). We found that the change in the β-MyHC/α-MyHC ratio was also evident at the RNA level, suggesting that lncMYH7b is affecting β-MyHC pre-translationally (Fig. 5A). To gain functional insight into the role of lncMYH7b in regulating cardiac gene expression, we performed a KEGG pathway analysis of our RNA-seq dataset and found that pathways associated with cardiomyopathies were affected by the reduction in lncMYH7b expression (Fig. 5D). This is consistent with the increases in lncMYH7b levels observed in tissue from patients with chronic heart disease (Fig S1). Of interest, expression of two members of the TEA domain (TEAD) transcription factor family, TEAD1 and TEAD3, was decreased in our lncMYH7b KD dataset. TEAD family members are known to bind to and enhance transcription at the β-MyHC promoter and are upregulated in hypertrophic cardiomyopathy patient hearts (25). TEAD1 knockout causes embryonic lethality in mice, while TEAD3 is less well studied but has a cardiac-enriched expression pattern (26). TEAD family members can complex with MEF2 transcription factor family members, which are master regulators of muscle formation and are known to enhance β-MyHC transcription (27–29). Moreover, further interrogation of the dataset revealed a significant decrease in the expression of MEF2A and PTK2, also known as focal adhesion kinase (FAK). FAK can complex with MEF2 in cardiomyocytes to enhance its activity under stress conditions, and in a mouse model where FAK was knocked out in cardiomyocytes, MEF2A protein levels were downregulated (30–32). Additionally, the FAK pathway is one of the top 5 enriched pathways in our KEGG analysis (Fig. 5D). Finally, FAK has also been identified as an upstream activator of TEAD transcriptional networks (33, 34). This, along with lncMYH7b’s cytoplasmic localization suggests that lncMYH7b is targeting an upstream effector of TEAD transcription in cardiomyocytes.

**Figure 5.**
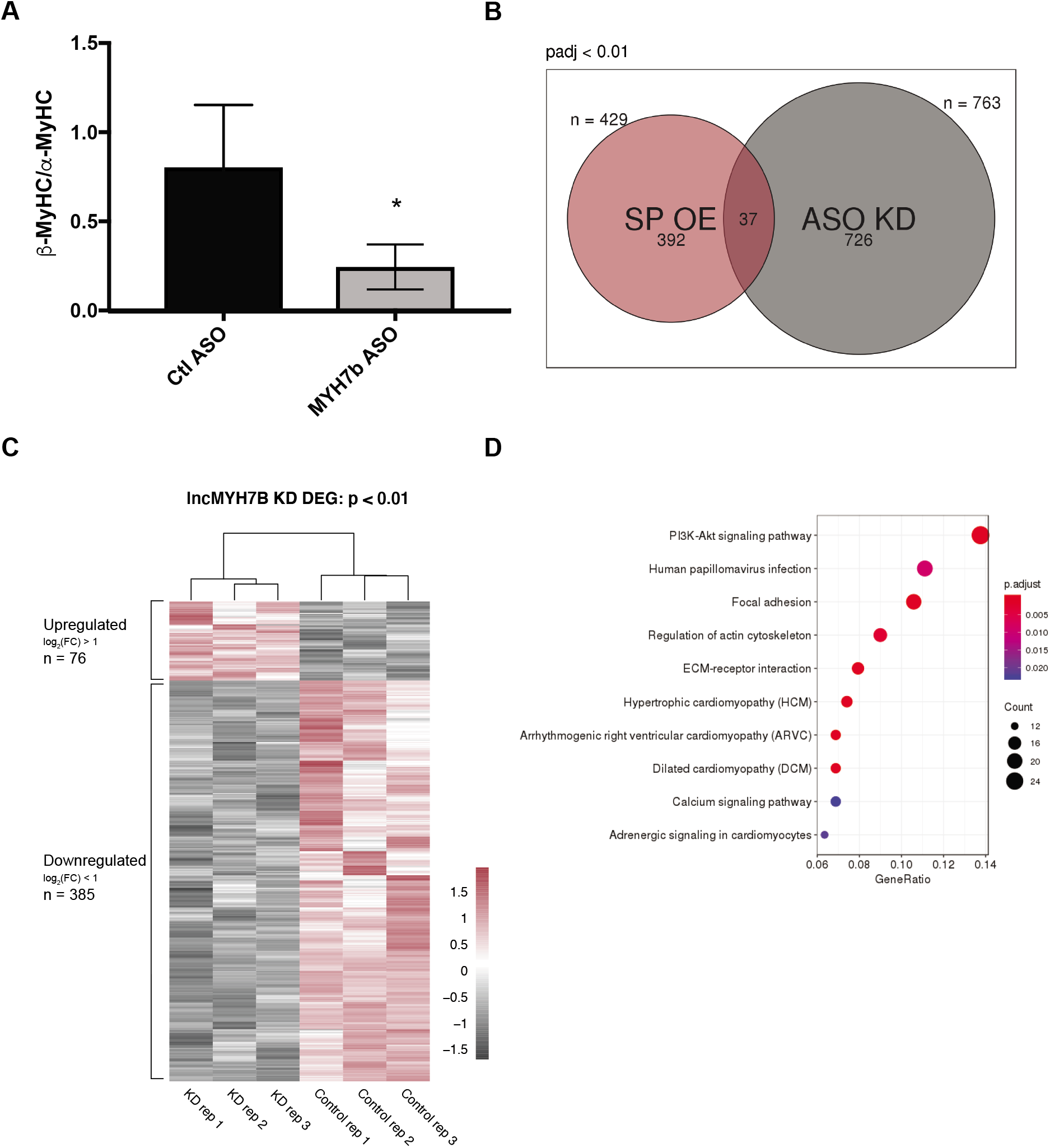
lncMYH7b KD shows little overlap with MYH7b_sp overexpression. **A)** RNA-seq of lncMYH7b KD in iPS-CMs shows a reduction in the β-/α-MyHC ratio, confirming that lncMYH7b is affecting β-MyHC RNA. **B)** Venn diagram of the differentially expressed genes from the MYH7b_sp overexpression and lncMYH7b KD shows that there is little overlap, which further supports the conclusion that MYH7b_sp is non-consequential. **C)** RNA-seq of lncMYH7b KD shows an enrichment of downregulated genes. **D)** KEGG analysis of the lncMYH7b KD RNA-seq shows enrichment of focal adhesion pathways, indicating possible role of FAK in lncMYH7b regulation.

## DISCUSSION

We have discovered that the MYH7b gene not only encodes a myosin heavy chain protein and a miRNA but also a regulatory lncRNA we have named lncMYH7b. This is the first example of a myosin heavy chain sense transcript with a regulatory role. Since MYH7b protein is undetectable in the mammalian heart despite the presence of the RNA, we postulated that the locus is transcribed to enable miR-499 expression. However, miR-499 null mice have no obvious cardiac phenotypes either at baseline or when stressed (13). Importantly, MYH7b RNA levels are preserved in those animals. Although it is a complicated locus encoding several potential active molecules, we present evidence that it is the lncRNA itself that has a regulatory role in cardiac gene expression.

There have been several reports of other lncRNAs having important roles in the heart. Myheart, the antisense transcript of β-MyHC, and Chaer are two examples of lncRNAs that regulate genes associated with heart disease (35, 36). However, most known cardiac lncRNAs work at the chromatin level by affecting epigenetic factors. lncMYH7b’s cytoplasmic localization implies a novel post-transcriptional mechanism of regulating gene expression (37, 38). There are examples of annotated lncRNAs encoding micropeptides, but despite *MYH7b* generating full-length protein in other tissues, there is no evidence for any peptide derived from this locus in the human heart (9, 11, 12, 18, 21, 22). Furthermore, this is an exciting example of the repurposing of an ancient proteincoding gene to produce a regulatory molecule via alternative splicing. The non-coding portion of the genome has been an area of exciting discovery in recent years, and the prospect of more bifunctional RNAs opens up even more possibilities.

This lncMYH7b-dependence of β-MyHC expression at both the RNA and protein levels is an exciting new discovery in the field of cardiac biology. An increase in the β-MyHC/α-MyHC ratio is a hallmark of heart disease (2, 3). This change in myosin heavy chain composition has been shown to compromise the contractility of cardiomyocytes. Heart failure patients that respond functionally to β-adrenergic receptor blocker treatment have a shift back towards the physiological β-MyHC/α-MyHC ratio (4). Therefore, understanding the regulation of β-MyHC expression is of great interest to the cardiac field. Since our FISH data show that lncMYH7b is primarily localized in the cytoplasm (Fig 3B), we do not think that lncMYH7b acts directly to regulate transcription from the β-MyHC locus. We analyzed the transcriptome of lncMYH7b KD hiPS-CMs to gain insight into potential networks that include this lncRNA. Notably, we found decreased expression of the transcription factors TEAD1, TEAD3, and MEF2A upon lncMYH7b KD. As these factors are known to transcriptionally regulate β-MyHC via a distal MCAT element, we suspect that this is a main driver for the observed decrease in the β-MyHC/α-MyHC ratio (25, 27). How lncMYH7b mediates this change in TEAD and MEF2 family member levels is an area of ongoing investigation. However, it is possible that the expression level changes in these transcription factors are downstream effects of changes in FAK levels. FAK not only affects the levels of MEF2A, but it is also a known upstream regulator of TEAD-mediated transcription(33, 39). All of these factors (FAK, MEF2, and TEADs) have major roles in both normal heart function and response to disease states, indicating that lncMYH7b may also have a role in physiologic cardiac gene expression regulation as well as a role in disease response. Further dissection of this regulatory network is necessary to better understand lncMYH7b’s role in the transcriptional landscape of the heart. Once fully defined, this offers multiple new target points for therapeutics to treat chronic heart disease.

## DATA AVAILABILITY

The RNA-seq datasets generated for this study are available on NCBI GEO GSE157260 (MYH7b_sp overexpression) and GSE157217 (KD study).

## Acknowledgments

We are grateful to Dr. Michael Bristow for helpful discussions.

## Funding and additional information

This work was primarily funded by NIH R01 GM029090 (to LAL), the NIH-CU Biophysics Training Grant (to LJB) and NIH NRSA F31HL147487 (to LJB).

## References

1. Weiss, A., and Leinwand, L. (1996) the Mammalian Myosin Heavy Chain Gene Family. Annu. Rev. Cell Dev. Biol. 12, 417–439

2. Nakao, K., Minobe, W., Roden, R., Bristow, M. R., and Leinwand, L. A. (1997) Myosin heavy chain gene expression in human heart failure. J. Clin. Invest. 100, 2362–2370

3. Miyata, S., Minobe, W., Bristow, M. R., and Leinwand, L. A. (2000) Myosin heavy chain isoform expression in the failing and nonfailing human heart. Circ. Res. 86, 386–390

4. Lowes, B. D., Gilbert, E. M., Abraham, W. T., Minobe, W. A., Larrabee, P., Ferguson, D., Wolfel, E. E., Lindenfeld, J. A., Tsvetkova, T., Robertson, A. D., Quaife, R. A., and Bristow, M. R. (2002) Myocardial gene expression in dilated cardiomyopathy treated with beta-blocking agents. N. Engl. J. Med. 346, 1357–1365

5. Hasegawa, K., Lee, S. J., Jobe, S. M., Markham, B. E., and Kitsis, R. N. (1997) cis-Acting sequences that mediate induction of beta-myosin heavy chain gene expression during left ventricular hypertrophy due to aortic constriction. Circulation. 96, 3943–53

6. Krenz, M., and Robbins, J. (2004) Impact of beta-myosin heavy chain expression on cardiac function during stress. J. Am. Coll. Cardiol. 44, 2390–2397

7. Rundell, V. L. M., Manaves, V., Martin, A. F., and de Tombe, P. P. (2005) Impact of beta-myosin heavy chain isoform expression on cross-bridge cycling kinetics. Am J Physiol Hear. Circ Physiol. 288, H896–903

8. Desjardins, P. R., Burkman, J. M., Shrager, J. B., Allmond, L. A., and Stedman, H. H. (2002) Evolutionary implications of three novel members of the human sarcomeric myosin heavy chain gene family. Mol. Biol. Evol. 19, 375–393

9. Lee, L. A., Karabina, A., Broadwell, L. J., and Leinwand, L. A. (2019) The ancient sarcomeric myosins found in specialized muscles. Skelet. Muscle. 9, 1–15

10. Bell, M. L., Buvoli, M., and Leinwand, L. A. (2010) Uncoupling of Expression of an Intronic MicroRNA and Its Myosin Host Gene by Exon Skipping. Mol. Cell. Biol. 30, 1937–1945

11. Rossi, A. C., Mammucari, C., Argentini, C., Reggiani, C., and Schiaffino, S. (2010) Two novel/ancient myosins in mammalian skeletal muscles: MYH14/7b and MYH15 are expressed in extraocular muscles and muscle spindles. J. Physiol. 588, 353–364

12. Rubio, M. D., Johnson, R., Miller, C. A., and Huganir, R. L. (2011) Regulation of Synapse Structure and Function by Distinct Myosin II Motors. J Neurosci. 31, 1448–1460

13. van Rooij, E., Quiat, D., Johnson, B. A., Sutherland, L. B., Qi, X., Richardson, J. A., Kelm, R. J., and Olson, E. N. (2009) A Family of microRNAs Encoded by Myosin Genes Governs Myosin Expression and Muscle Performance. Dev. Cell. 17, 662–673

14. Sadayappan, S., Gulick, J., Klevitsky, R., and Lorenz, J. N. (2010) Cardiac Myosin Binding Protein-C Phosphorylation in a “Humanized” Mouse Heart. Physiology. 119, 1253–1262

15. Lian, X., Hsiao, C., Wilson, G., Zhu, K., Hazeltine, L. B., Azarin, S. M., Raval, K. K., Zhang, J., Kamp, T. J., and Palecek, S. P. (2012) Robust cardiomyocyte differentiation from human pluripotent stem cells via temporal modulation of canonical Wnt signaling. Proc. Natl. Acad. Sci. U. S. A. 10.1073/pnas.1200250109

16. Nag, S., Sommese, R. F., Ujfalusi, Z., Combs, A., Langer, S., Sutton, S., Leinwand, L. A., Geeves, M. A., Ruppel, K. M., and Spudich, J. A. (2015) Contractility parameters of human b-cardiac myosin with the hypertrophic cardiomyopathy mutation R403Q show loss of motor function. Sci. Adv. 1, 1–16

17. Peter, A. K., Rossi, A. C., Buvoli, M., Ozeroff, C. D., Crocini, C., Perry, A. R., Buvoli, A. E., Lee, L. A., and Leinwand, L. A. (2019) Expression of Normally Repressed Myosin Heavy Chain 7b in the Mammalian Heart Induces Dilated Cardiomyopathy. J. Am. Heart Assoc. 10.1161/JAHA.119.013318

18. Haraksingh, R. R., Jahanbani, F., Rodriguez-Paris, J., Gelernter, J., Nadeau, K. C., Oghalai, J. S., Schrijver, I., and Snyder, M. P. (2014) Exome sequencing and genome-wide copy number variant mapping reveal novel associations with sensorineural hereditary hearing loss. BMC Genomics. 15, 1155

19. Chistiakov, D. A., Orekhov, A. N., and Bobryshev, Y. V. (2016) Cardiac-specific miRNA in cardiogenesis, heart function, and cardiac pathology (with focus on myocardial infarction). J. Mol. Cell. Cardiol. 94, 107–121

20. Trcek, T., Sato, H., Singer, R. H., and Maquat, L. E. (2013) Temporal and spatial characterization of nonsense-mediated mRNA decay. Genes Dev. 27, 541–551

21. Nelson, B. R., Makarewich, C. A., Anderson, D. M., Winders, B. R., Troupes, C. D., Wu, F., Reese, A. L., McAnally, J. R., Chen, X., Kavalali, E. T., Cannon, S. C., Houser, S. R., Bassel-duby, R., and Olson, E. N. (2016) A peptide encoded by a transcript annotated as long noncoding RNA enhances SERCA activity in muscle. Science (80-.). 351, 271–275

22. Anderson, D. M., Anderson, K. M., Chang, C. L., Makarewich, C. A., Nelson, B. R., McAnally, J. R., Kasaragod, P., Shelton, J. M., Liou, J., Bassel-Duby, R., and Olson, E. N. (2015) A micropeptide encoded by a putative long noncoding RNA regulates muscle performance. Cell. 160, 595–606

23. Zhao, H., Dahlö, M., Isaksson, A., Syvänen, A. C., and Pettersson, U. (2012) The transcriptome of the adenovirus infected cell. Virology. 424, 115–128

24. Ewels, P. A., Peltzer, A., Fillinger, S., Alneberg, J., Patel, H., Wilm, A., Garcia, M. U., Tommaso, P. Di, and Nahnsen, S. (2019) nf-core: Community curated bioinformatics pipelines. bioRxiv. 10.1101/610741

25. Iwaki, H., Sasaki, S., Matsushita, A., Ohba, K., and Matsunaga, H. (2014) Essential Role of TEA Domain Transcription Factors in the Negative Regulation of the MYH 7 Gene by Thyroid Hormone and Its Receptors. PLoS One. 10.1371/journal.pone.0088610

26. Jin, Y., Messmer-Blust, A. F., and Li, J. (2011) The Role of Transcription Enhancer Factors in Cardiovascular Biology. Trends Cardiovasc. Med. 21, 1–5

27. Karasseva, N., Tsika, G., Ji, J., Zhang, A., Mao, X., and Tsika, R. (2003) Transcription Enhancer Factor 1 Binds Multiple Muscle MEF2 and A/T-Rich Elements during Fast-to-Slow Skeletal Muscle Fiber Type Transitions. Mol. Cell. Biol. 23, 5143–5164

28. Wales, S., Hashemi, S., Blais, A., and McDermott, J. C. (2014) Global MEF2 target gene analysis in cardiac and skeletal muscle reveals novel regulation of DUSP6 by p38MAPK-MEF2 signaling. Nucleic Acids Res. 42, 11349–11362

29. Potthoff, M. J., and Olson, E. N. (2007) MEF2: a central regulator of diverse developmental programs. Development. 134, 4131–4140

30. Cardoso, A. C., Pereira, A. H. M., Ambrosio, A. L. B., Consonni, S. R., Rocha de Oliveira, R., Bajgelman, M. C., Dias, S. M. G., and Franchini, K. G. (2016) FAK Forms a Complex with MEF2 to Couple Biomechanical Signaling to Transcription in Cardiomyocytes. Structure. 24, 1301–1310

31. Nadruz, W., Corat, M. A. F., Marin, T. M., Guimarães Pereira, G. A., and Franchini, K. G. (2005) Focal adhesion kinase mediates MEF2 and c-Jun activation by stretch: Role in the activation of the cardiac hypertrophic genetic program. Cardiovasc. Res. 68, 87–97

32. Peng, X., Wu, X., Druso, J. E., Wei, H., Park, A. Y. J., Kraus, M. S., Alcaraz, A., Chen, J., Chien, S., Cerione, R. A., and Guan, J. L. (2008) Cardiac developmental defects and eccentric right ventricular hypertrophy in cardiomyocyte focal adhesion kinase (FAK) conditional knockout mice. Proc. Natl. Acad. Sci. U. S. A. 105, 6638–6643

33. Kim, N. G., and Gumbiner, B. M. (2015) Adhesion to fibronectin regulates Hippo signaling via the FAK-Src-PI3K pathway. J. Cell Biol. 210, 503–515

34. Park, J. H., Shin, J. E., and Park, H. W. (2018) The role of hippo pathway in cancer stem cell biology. Mol. Cells. 41, 83–92

35. Han, P., and Chang, C. P. (2015) Myheart hits the core of chromatin. Cell Cycle. 14, 787–788

36. Wang, Z., Zhang, X., Ji, Y., Zhang, P., Yokota, T., Ang, Y. S., Li, S., Cass, A., and Vondriska, T. M. (2017) A Long Non-Coding RNA Defines an Epigenetic Checkpoint in Cardiac Hypertrophy. Nat. Med. 22, 1131–1139

37. Chang, C. P., and Han, P. (2016) Epigenetic and lncRNA regulation of cardiac pathophysiology. Biochim. Biophys. Acta - Mol. Cell Res. 1863, 1767–1771

38. Rashid, F., Shah, A., and Shan, G. (2016) Long Non-coding RNAs in the Cytoplasm. Genomics, Proteomics Bioinforma. 14, 73–80

39. Torsoni, A. S., Marin, T. M., Velloso, L. A., and Franchini, K. G. (2005) RhoA/ROCK signaling is critical to FAK activation by cyclic stretch in cardiac myocytes. Am. J. Physiol. Circ. Physiol. 289, H1488–H1496

